# Influence of cognitive control on semantic representation

**DOI:** 10.1101/067553

**Authors:** Waitsang Keung, Daniel Osherson, Jonathan D. Cohen

**Affiliations:** Princeton Neuroscience Institute, Princeton University, NJ 08540, US; Department of Psychology, Princeton University, NJ 08540, US

**Author notes:** Corresponding author: Princeton Neuroscience Institute, Princeton University, Princeton, NJ 08540, US.

## Abstract

The neural representation of an object can change depending on its context. For instance, a horse may be more similar to a bear than to a dog in terms of size, but more similar to a dog in terms of domesticity. We used behavioral measures of similarity together with representational similarity analysis and functional connectivity of fMRI data in humans to reveal how the neural representation of semantic knowledge can change to match the current goal demand. Here we present evidence that objects similar to each other in a given context are also represented more similarly in the brain and that these similarity relationships are modulated by context specific activations in frontal areas.

**Significance statement:** The judgment of similarity between two objects can differ in different contexts. Here we report a study that tested the hypothesis that brain areas associated with task context and cognitive control modulate semantic representations of objects in a task-specific way.

We first demonstrate that task instructions impact how objects are represented in the brain. We then show that the expression of these representations is correlated with activity in regions of frontal cortex widely thought to represent context, attention and control.

In addition, we introduce spatial variance as a novel index of representational expression and attentional modulation. This promises to lay the groundwork for more exacting studies of the neural basis of semantics, as well as the dynamics of attentional modulation.

## Introduction

People often view the same set of stimuli differently under different contexts. For example, a horse may be more similar to a bear than to a dog in terms of size, but more similar to a dog in terms of domesticity. Similarity relationships among a given set of objects need not be fixed; they can change to match the current context or task demand. Evidence (Çukur et al., 2013) suggest that similarity among object representations changes according to task demand. But it is not yet clear what neural mechanism effects such a change in representation.

Previous theories have proposed that brain areas known to be involved in cognitive control guide the flow of information in posterior cortices to execute task relevant behaviors (Miller and Cohen, 2001). Such cognitive control might effect shifts in attention to distinct dimensions of stimuli, thereby affecting how they are represented (Smith et al., 1974; Cohen et al., 1990; Kanwisher and Wojciulik, 2000; Maunsell and Treue, 2006), and thus, how similar they seem to be. We hypothesized that task demands affect semantic representations of objects and thereby similarity judgments about these objects, and that this effect is modulated in a task-specific way by activations in the frontal brain areas.

Category membership of objects has been reliably decoded in functional magnetic resonance imaging (fMRI) studies using both univariate (Kanwisher and McDermott, 1997; McCarthy et al., 1997; Aguirre et al., 1998; Epstein and Kanwisher, 1998; Chao et al., 1999; Gauthier et al., 1999; 2000) and multivariate methods (Haxby et al., 2001; Cox and Savoy, 2003; Kay et al., 2008; Haxby et al., 2011; Kriegeskorte, 2011; Connolly et al., 2012). Multivariate pattern analysis methods have been especially useful in extracting and characterizing fine-grained information carried by fMRI data. One such method is *representational similarity analysis* (RSA) (Kriegeskorte et al., 2008). RSA explores the similarity structure among neural representations and provides a means to relate neural data to behavior (Kriegeskorte, 2011; Kriegeskorte and Kievit, 2013). Previous studies have shown that objects’ similarity relationships can be extracted from human and monkey neural data using RSA (Hanson et al., 2004; O’Toole et al., 2007; Kriegeskorte et al., 2008; Weber et al., 2009; Bruffaerts et al., 2013). Several studies using pictures have suggested that neural representations of visual images in human participants emphasize categorical boundaries, i.e. representations of images that belong in the same category tend to form clusters in representational space (Kriegeskorte and Kievit, 2013). More importantly, recent studies have tried to disentangle the effects of perceptual factors in visual images of objects from conceptual factors in the concepts of objects by using words instead of pictures as stimuli (Bruffaerts et al., 2013). These studies were able to decode the category of the objects or extract the similarity relationships among objects from neural data. In this study, we utilized representational similarity analysis to uncover brain areas that best represent context-specific semantic representation.

To test our hypotheses, we asked participants to perform a similarity comparison task on a set of animals and, in a separate study, fruits. Before scanning, we obtained behavioral ranking of the similarity among the items (animals or fruits) under two different semantic contexts (domesticity and size for animals, taste and size for fruits). Then, while being scanned using fMRI, we asked the participants to compare the items under each of the two different semantic contexts. We expected to find brain areas for which the similarity relationship among neural representations of the items correlated with that of the participant’s behavioral judgment of similarity. Importantly, we predicted this correlation to be context-specific; that is, we expected that the neural similarity relationship among the items would change according to the context under which they were being compared, so as to match the behavioral similarity judgment under that context.

To identify areas the activity of which might be related to the expression of context-specific similarity structure (and thus are candidates for the source of dimension-specific context effects), we performed a functional connectivity analysis. Using areas identified in the foregoing similarity analysis as seed regions, we examined correlations between these areas and the rest of the brain separately under the two different contexts. Notably, we correlated the spatial variance (variance across voxels at each time point) of the seed regions with the activation of the rest of the brain, using spatial variance as a proxy for the strength of representation. Our rationale was that, if attention acts to modulate representations (e.g. by influencing the gain of activity within the target region (Cohen et al., 1990; McAdams and Maunsell, 1999; Treue and Trujillo, 1999; Reynolds et al., 2000; Maunsell and Treue, 2006; Aboitiz and Cosmelli, 2008)), then increased attention should increase the activity of activated voxels and decrease the activity of suppressed voxels, thus increasing the variance of activity across voxels within the region. Thus, we used spatial variance as an estimate of gain within a region; and we sought to identify areas the activity of which was correlated with such changes in gain, as candidates for the source of context (control) signals responsible for the changes in gain. We found significant correlations between neural and behavioral similarity for domesticity but not for size in animals. We also found significant correlations for taste but not for size in fruits. Our connectivity analysis revealed correlations between regions that reflected behavioral similarity and other brain areas including dorsolateral prefrontal cortex (dlPFC) and inferior frontal gyrus (IFG) that were not only significant within context but also significantly differentiated between the two contexts. We also compared the results against that of using activation only connectivity analysis and found significant results only in the analysis using spatial variance, consistent with our hypothesis about the mechanism of action of attention (modulation rather than a change in mean activity). Together, these results suggest that the neural representation of objects reflects behaviorally determined, context-specific similarity structure, and that this may be expressed as a change in the gain of their representation (i.e., contrast) rather than a change in mean activity. They also suggest that the change in neural representation is modulated by context-specific activity in regions of frontal cortex.

## Methods

### Participants

Twenty-four participants (between 18-26 years old, fifteen women) performed the main fMRI study with animal stimuli. Sixteen participants (between 18-26 years old, thirteen women) performed the replication fMRI study with fruit stimuli. All experimental procedures were approved by the Institutional Review Board at Princeton University. Written informed consent was obtained from all participants.

### Stimuli and experimental procedure

Stimuli for the main fMRI study comprised twelve animals that varied across two dimensions: domesticity and size (Figure 1a). Each dimension had two features (“wild” and “domestic” for domesticity, “big” and “small” for size). Before the fMRI experiment, participants first ranked the animals along the two dimensions, from domestic to wild for domesticity and small to big for size.

**Figure 1.**
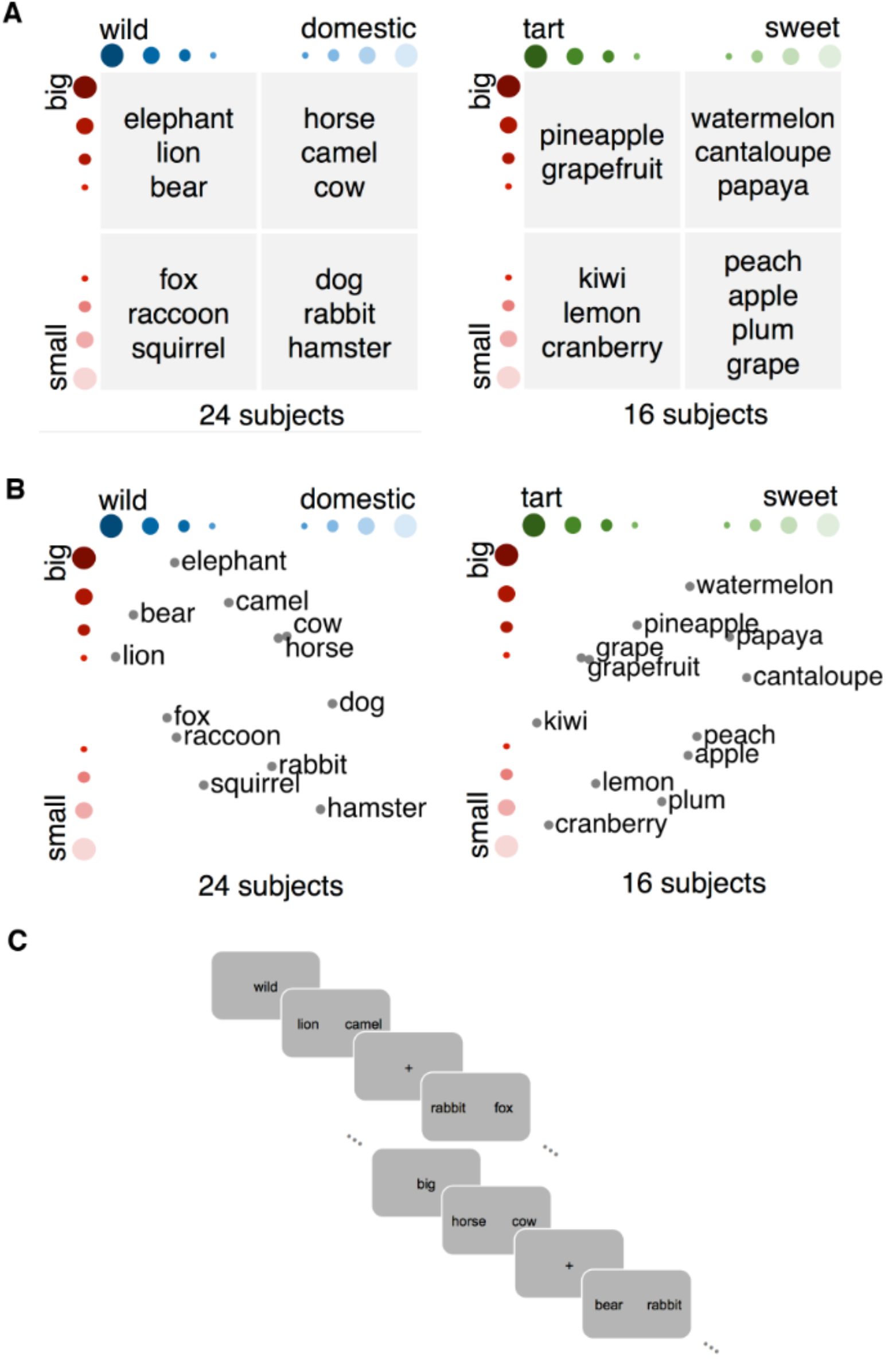
Design, stimuli and behavioral results. (a) Animal stimuli that vary along domesticity and size. Fruits stimuli that vary along taste and size. (b) Scatter plot of averaged participants’ behavioral ranking of stimuli across two dimensions. Individual participant’s behavioral ranking was used in subsequent analyses. (c) Schematic of the fMRI experimental design.

In the scanner, participants compared between pairs of stimuli presented in an event-related design (Figure 1c). At the beginning of each block, a cue was presented for 4 s instructing the participants which feature to pay attention to. The cue was either “wild”, “domestic”, “big” or “small”. In each trial, participants were shown a pair of animals positioned on the left and right sides of the screen in the form of words in black against a grey background and participants then decided which animal of the pair was best described by the cued feature. For example, if the cue was “big”, they needed to decide which animal was bigger. The pair of animals stayed on the screen for 2 s, after which a fixation cross appeared on the screen. Participants had to make a button press response within 1 s after the fixation cross appeared. Intertrial intervals were jittered between 4 – 8s. Participants in the fruits experiment made comparisons among twelve fruits along the dimensions taste (“sweet” and “tart”) and size.

All possible pairs of twelve animals were presented once under each feature. There were a total of eleven scanning runs, and four blocks for all four features in each run. The order of features in each run was randomized. Each block had six trials. This behavioral ranking was used to gauge their performance in the scanner and to generate the experiment in such a way that left and right button presses were balanced in each block. Stimuli were presented using MATLAB software (MathWorks) and the Psychophysics Toolbox using a projector outside the MRI scanner that displayed the stimuli onto a translucent screen located at the end of the scanner bore, which participants viewed through a mirror attached to the head coil.

### fMRI data acquisition and preprocessing

Functional (EPI sequence, 34 slices with full cerebrum coverage, resolution = 3 × 3 × 3 mm with 1 mm gap, repetition time = 2.0 s, echo time = 30 ms, flip angle = 90°) and anatomical (MPRAGE sequence, 256 matrix, repetition time = 25 s, echo time = 438 ms, flip angle = 8°, resolution = 1 × 1 × 1 mm) images were acquired using a 3T Skyra MRI scanner (Siemens) at Princeton University. Data were processed using MATLAB and SPM8 (Wellcome Trust Centre for Neuroimaging, University College London). Functional data were motion corrected, and low-frequency drifts were removed with a temporal high-pass filter (cutoff of 0.0078 Hz). Images were normalized to Montreal Neurological Institute (MNI) coordinates. Images were spatially smoothed with a FWHM 5 kernel. In addition to the standard fMRI preprocessing procedure, reaction time was used as a regressor and was regressed out of the data.

### Univariate contrasts

The main purpose of the univariate contrast was to find each animal’s neural representation as an input into generating the neural similarity matrix. A total of 24 General Linear Model (GLM) analyses were run using SPM, one for each animal under each dimension (12 animals × 2 dimensions). In each GLM, two regressors were set up: a regressor that indicated the trials in which a certain animal appeared under a certain dimension, and another regressor that indicated all the trials in the opposite dimension. These indicators were convolved with a standard hemodynamic response function. The representation of a certain animal under a certain dimension was defined by the contrast of trials in which the animal appeared in that dimension and all other trials in the opposite dimension. We hoped to average out the representation of other animals with GLM. For balance, all pairings for a certain animal appeared the same number of times under one dimension vs. the other dimension.

### Similarity searchlight analysis

A searchlight analysis was performed across the whole brain to find areas that best predicted behavioral data. For each participant, a similarity matrix among twelve animals was generated for each dimension based on his/her behavioral ranking of the animals under that dimension. We used a searchlight sphere of 4 voxels radius. For each searchlight sphere, a similarity matrix was computed from the neural representations of the animals (generated from the previous univariate contrasts) for each dimension. We then correlated the two similarity matrices from behavioral ranking and from neural representations and got a correlation coefficient for that sphere. The highest correlation coefficient across the whole brain was selected for each participant. To make sure that it is dimension-specific, we used the exact same sphere and correlated its neural similarity matrix with the behavioral similarity matrix of the opposite dimension. We took the absolute value of this correlation and subtracted it from the original correlation coefficient (Fisher transformed). This way, we made sure that the neural data under one dimension was not predictive of behavioral data under the other.

To analyze the data statistically, we used permutation test in which we scrambled the behavioral rankings and repeated the analysis to determine a null distribution and a p value for each voxel. At the group level, we then computed, for each voxel, the number of participants that passed the p = 0.05 threshold, and performed a binomial test. To correct for multiple comparisons, we thresholded the group result with p = 0.05 from the binomial test, and used AlphaSim (AFNI) to perform a randomization test on the group data to threshold the cluster size.

Separately, we also performed a group-level statistical test on the maximum correlation coefficient in each participant. We corrected for multiple comparisons by using a permutation test: we scrambled the behavioral ranking and repeated the analysis, and repeated this 1000 times. This generated a null distribution of the dimension-specific correlation values for each participant. We then identified searchlights for which the dimension-specific correlation value passed p = 0.05 in the null distribution. Finally, we submitted the number of participants for whom one or more searchlights were significant to a binomial test for group level statistics. In an additional test, we averaged the dimension-specific correlation values across all participants for each dimension. To determine whether this group averaged maximum correlation coefficient was significant, we again did a permutation test and generated a null distribution of group averaged maximum correlation coefficients. The real group average was compared to this distribution to determine the p-value.

### Connectivity searchlight analysis

To determine whether the areas expressing similarity representations interacted with other areas in the brain, we used spheres in which neural similarity best correlated with behavioral similarity as seed regions to compute the correlation between these regions and the rest of the brain for each individual. Specifically, we computed the correlation between the spatial variance time series of the seed regions and the activation time series of the rest of the brain. To create the spatial variance time series, for each time point, we computed the variance across voxels within the searchlight sphere (Figure 3b). The purpose was to use spatial variance as a measure of the strength of similarity representation time point to time point. To make sure that the correlation is dimension-specific, we computed correlations for two time series respectively for domesticity trials and for size trials, and contrasted the two correlations (Figure 3a). For example, to look at the correlation between seed region that showed the best similarity representation of animals in domesticity and the rest of the brain, we computed the correlation between these two areas during size trials and subtracted from it the correlation during domesticity trials.

**Figure 2.**
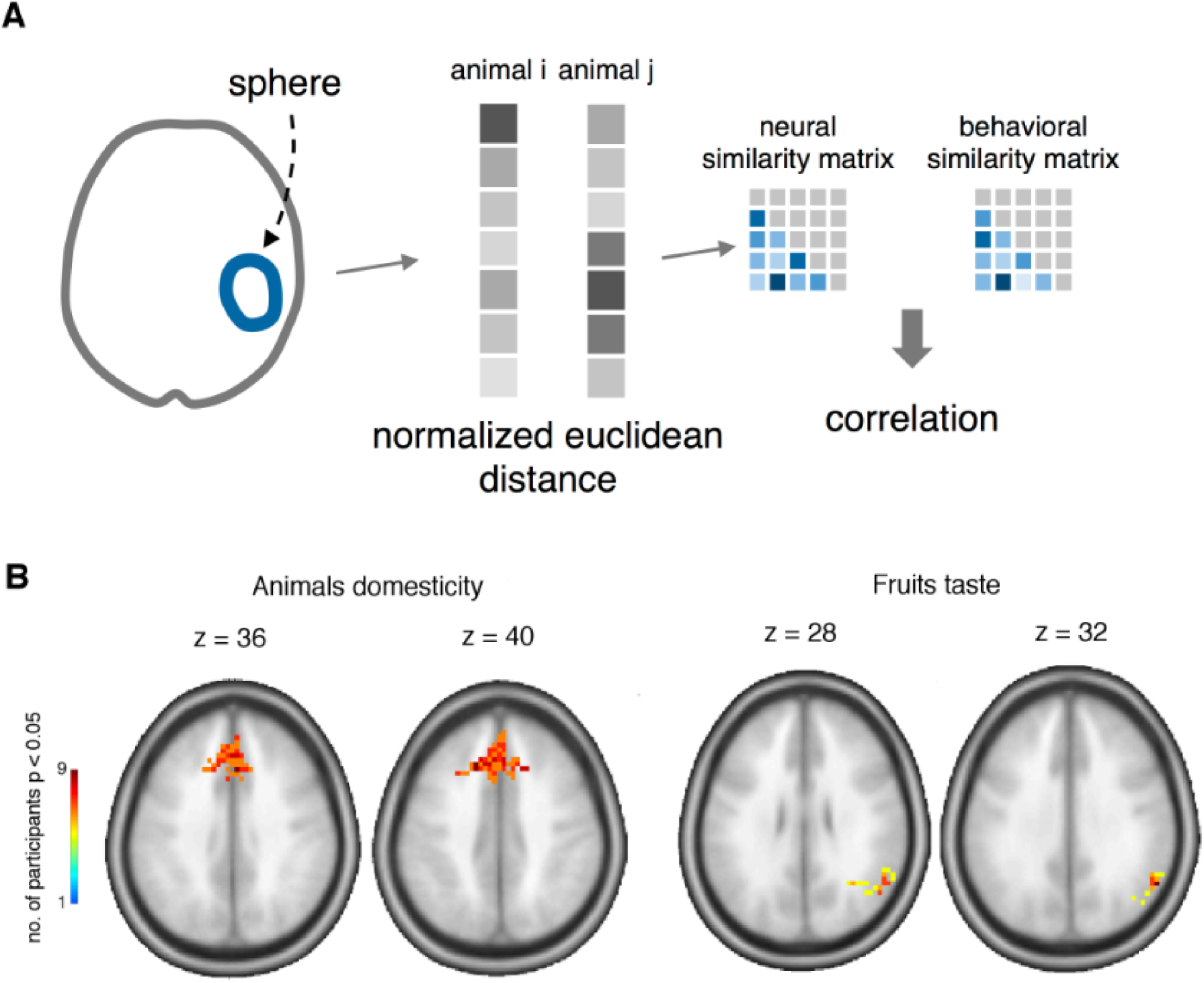
Similarity analysis. (a) Schematic of similarity searchlight analysis. Neural representation for each animal was extracted in GLM analysis, and was used to construct a neural similarity matrix to predict similarity matrix constructed from behavioral ranking data. (b) Similarity analysis results. Cluster that spans MFG and dACC reliably predict dimension-specific behavioral judgment of similarity for animals domesticity, while as cluster in IPS reliably predict that for fruits taste.

**Figure 3.**
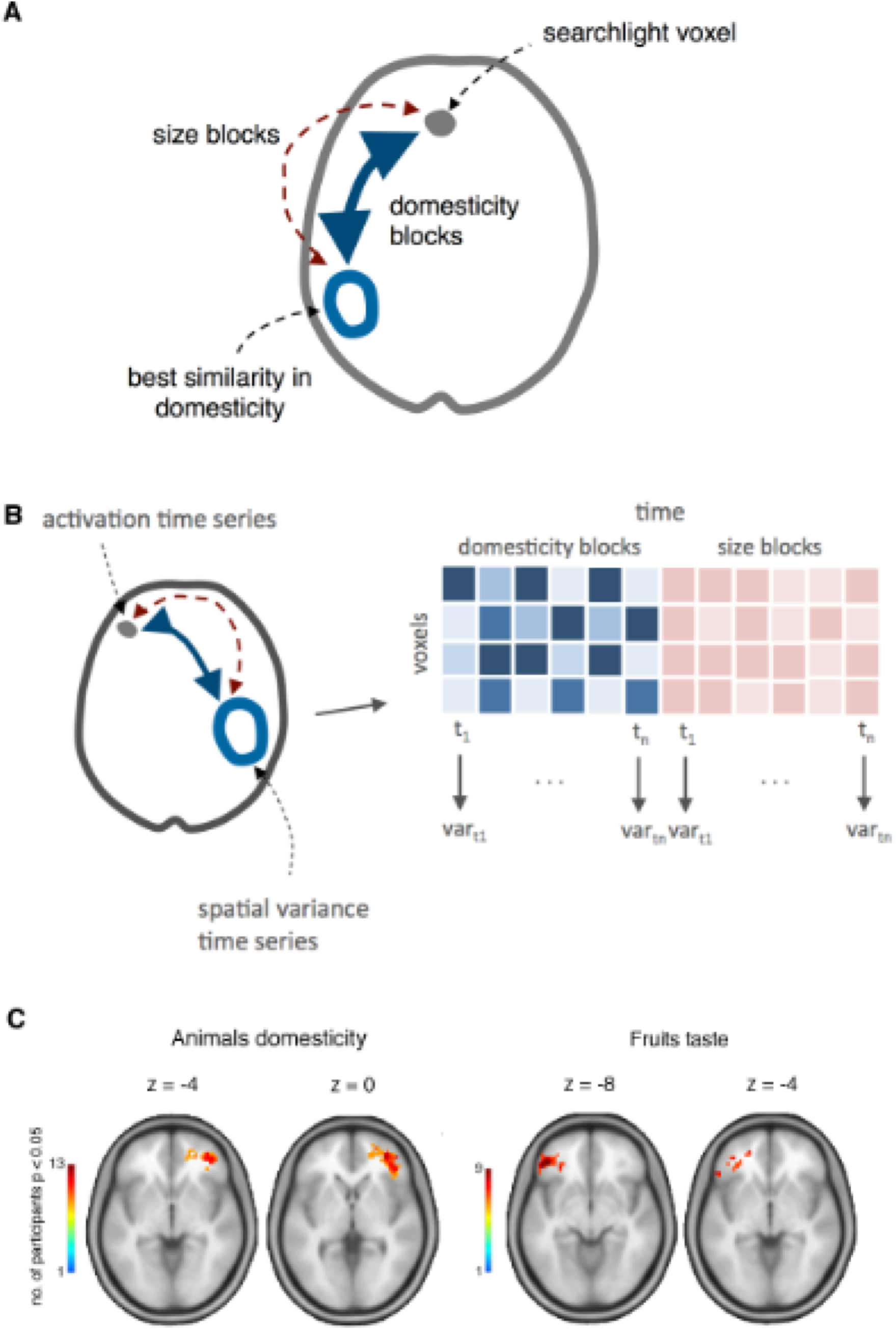
Connectivity analysis. (a-b) Schematic of connectivity analysis. (a) Areas that best predicted behavioral similarity were used as seed regions. Correlation between seed regions and other parts of the brain during trials of one dimension was compared against that of the other dimension. (b) Time series of spatial variance of seed region was generated by computing the variance across voxels at each given time point. (c) Connectivity analysis results. Clusters in right IFG and right dlPFC reliably predict dimension-specific changes in spatial variance in the seed regions in animals domesticity. Clusters in left IFG and dlPFC reliably predict dimension-specific changes in spatial variance in the seed regions in fruits taste.

Similar to the statistics done in the similarity searchlight analysis, we used permutation test in which we scrambled the time series of the seed regions and repeated the analysis to determine a null distribution and a p value for each voxel in each subject. At the group level, we then computed, for each voxel, the number of participants that passed the p = 0.05 threshold, and performed a binomial test. To correct for multiple comparisons, we thresholded the group result with p = 0.05 from the binomial test, and used AlphaSim (AFNI) to perform a randomization test on the group data to threshold the cluster size.

We also performed a group-level statistical test on the maximum correlation coefficient in each participant. We picked the maximum correlation coefficient for each participant under one dimension and subtracted from it the correlation between the same areas under the opposite dimension. We performed a permutation test in which we scrambled the activation time series 1000 times to compute a null distribution of the dimension-specific correlation values for each participant. We then identified searchlights for which the dimension-specific correlation value passed p = 0.05 in the null distribution. Finally, we submitted the number of participants for whom one or more searchlights were significant to a binomial test for group level statistics. We also did an additional test by averaging the dimension-specific correlation values across all participants for each dimension. We performed a permutation test to compute a null distribution of the group average of the maximum dimension-specific correlation. A p-value is obtained for each of the two dimensions.

## Results

### Behavioral data

In the first experiment, we used 12 animal stimuli that varied systematically along two dimensions: domesticity and size (Figure 1a). We repeated the experiment in a separate group of participants using fruits as stimuli instead, with taste and size as the two dimensions. Individual specific behavioral rankings were obtained by asking participants to rank the animals from “domestic” to “wild” for domesticity and “small” to “big” for size (“sweet” to “tart” for taste and “small” to “big” for size in fruits) before scanning. For each stimulus type (animals or fruits), the similarity matrices generated from the two dimensions were not significantly correlated with each other (animals r=0.027, t_23_=0.68, p=0.50; fruits r=0.058, t_23_=1.62, p=0.11). This ensured that the two dimensions were behaviorally distinguishable from one another, and thus had the greatest opportunity to yield dimension-specific neural representations.

### Similarity searchlight analysis

To test whether objects that were rated as similar to each other were also represented more similarly in the brain, we compared object similarity matrices generated from the behavioral data (outside the scanner) with similarity matrices generated from the participants’ fMRI data acquired while performing an object comparison task in the scanner.

In the scanner, at the beginning of each block of trials, participants were cued with an “anchoring feature” along one of the two dimensions (e.g., for animals: “domestic”, “wild”, “small” or “big”). On each trial of the block, they were shown a pair of stimuli and asked to decide which one of the two was best described by the cued feature. For example, in the animals experiment, if the cue was “big” participants needed to decide which animal was bigger (Figure 1c). We extracted the dimension-specific representation of each animal by running a Generalized Linear Model (GLM) with a regressor that indicated all the trials involving a specific animal judged along a specific dimension. Furthermore, to remove the potential confound of task difficulty as a factor in the neural similarity measurements, we used reaction time as proxy for task difficulty and regressed it out of the signal before analysis (Todd et al., 2013).

To examine the relationship of neural similarity to behavioral similarity along each dimension, we examined the dimension-specific neural representations over local searchlights throughout the entire brain. Within each participant, we compared the similarity matrix generated from the neural representations within a spotlight with that from that participant’s behavioral data (Figure 2a). We then identified searchlights that best predicted each participant’s behavioral similarity scores. To ensure that the neural similarity matrices were dimension-specific and did not predict behavioral data for the other dimension, we subtracted the absolute correlation between neural data for one dimension and behavioral data for the other dimension from the correlation between neural and behavioral data for the same dimension. We corrected for multiple comparisons by using a permutation test, by generating a null distribution of the dimension-specific correlation values for each participant, and then identifying searchlights for which the dimension-specific correlation value passed p = 0.05 in the null distribution. Finally, we submitted the number of participants for whom one or more searchlights were significant to a binomial test for group level statistics. For animals, we found a significant number of participants who exhibited searchlights exceeding p = 0.05 for domesticity (6 out of 24, binomial p (one-tailed) =10^-4^, average r = 0.54) across participants, but only a trending significance for size (3 out of 24, binomial p (one-tailed) = 0.086, average r = 0.51). For fruits, we found regions exhibiting a significant correlation for taste (6 out of 16, binomial p (one-tailed) = 10^-5^, average r = 0.57) but, again, not for size (2 out of 16, binomial p (one-tailed) = 0.15, average r = 0.51). In addition, we also compared the group average of the best correlations across subjects against the null distribution generated by the permutation test. We found the group average to be significant for domesticity in animals (p = 0.045) and trending significant for taste in fruits (p = 0.069). We did not find significant results for size in both animals (p = 0.8) and fruits (p = 0.47).

To identify the regions that exhibited significant effects in the similarity analysis at the group level, we used a permutation test to threshold the *p* value and a randomization test to threshold the cluster size. For domesticity in animals, the largest significant cluster across participants spanned middle frontal gyrus (MFG) and dACC (p < 0.05 corrected; Figure 2b; Table 2). For fruits, the largest significant cluster spanned right inferior parietal lobule (IPL) and right angular gyrus (AG) (p < 0.02 corrected; Figure 2b; Table 2) for taste.

**Table 1.** Seed regions for connectivity analysis - best performing searchlights in similarity analysis.

| Subj<br># | Regions | Brodmann<br>areas | x | y | z | Ext<br>ent |
| --- | --- | --- | --- | --- | --- | --- |
| Animals group Domesticity |  |  |  |  |  |  |
| 1 | Anterior cingulum | 24/32 | 3 | 35 | 6 | 202 |
| 2 | Postcentral gyrus | 2/3/4 | -33 | -34 | 70 | 153 |
| 3 | Inferior parietal lobule/ angular gyrus | 7/40 | -36 | -49 | 46 | 168 |
| 4 | Thalamus | 23 | 9 | -28 | 14 | 139 |
| 5 | Putamen/ caudate | 13 | -21 | 17 | 6 | 184 |
| 6 | Middle frontal gyrus | 10/46 | 39 | 44 | 22 | 152 |
| 7 | Middle frontal gyrus | 9 | 45 | 35 | 38 | 88 |
| 8 | Precuneus | 7 | 0 | -67 | 38 | 286 |
| 9 | Insula/ inferior frontal gyrus | 13/47 | 27 | 20 | -2 | 172 |
| 10 | Medial frontal gyrus | 10 | -9 | 50 | 26 | 175 |
| 11 | Medial frontal gyrus/ anterior cingulate | 9/10/32 | -3 | 47 | 14 | 279 |
| 12 | Medial frontal gyrus | 8/9 | 6 | 38 | 38 | 235 |
| 13 | Middle/ inferior frontal gyrus | 10/11/47 | -45 | 47 | -10 | 90 |
| 14 | Postcentral gyrus/ inferior parietal lobule | 5/3/40 | -30 | -43 | 58 | 183 |
| 15 | Inferior parietal lobule | 40 | 39 | -46 | 34 | 80 |
| 16 | Parahippocampal gyrus/ hippocampus | 20/28/34/3<br>6 | -24 | -10 | -30 | 110 |
| 17 | Inferior temporal gyrus/ temporal pole | 20/21/38 | -45 | -1 | -26 | 175 |
| 18 | Superior parietal lobule/ precuneus | 7 | -18 | -70 | 58 | 126 |
| 19 | Superior/ medial frontal gyrus | 10/11 | -15 | 56 | -14 | 71 |
| 20 | Middle frontal gyrus | 6/8/9 | -27 | 14 | 42 | 118 |
| 21 | Medial frontal gyrus/ cingulate gyrus | 9/32 | -15 | 29 | 34 | 143 |
| 22 | Middle frontal gyrus | 11/10 | -42 | 53 | -14 | 62 |
| 23 | Fusiform gyrus | 37 | -39 | -64 | -18 | 69 |
| 24 | Temporal pole | 38 | 30 | 17 | -30 | 115 |
| Fruits group Taste |  |  |  |  |  |  |
| 1 | Posterior cingulum | 7/23/31 | 0 | -49 | 30 | 257 |
| 2 | Thalamus | 23 | -9 | -19 | 2 | 224 |
| 3 | Inferior parietal lobule | 40 | -54 | -55 | 42 | 117 |
| 4 | Temporal pole | 38 | 42 | 14 | -30 | 193 |
| 5 | Insula/ Putamen | 13 | -33 | 5 | -6 | 192 |
| 6 | Superior frontal gyrus | 10 | -24 | 59 | 2 | 163 |
| 7 | Medial frontal gyrus | 11 | 0 | 56 | -14 | 146 |
| 8 | Inferior occipital gyrus | 18 | 36 | -88 | -14 | 104 |
| 9 | Inferior temporal gyrus/ inferior occipital gyrus | 19/37 | -48 | -70 | -14 | 124 |
| 10 | Middle frontal gyrus | 46 | -45 | 35 | 22 | 118 |
| 11 | Thalamus | 23 | -9 | -19 | 2 | 224 |
| 12 | Lingual gyrus | 19 | 21 | -58 | -6 | 138 |
| 13 | Temporal pole | 38 | -42 | 8 | -34 | 135 |
| 14 | Anterior cingulum | 10/32 | -9 | 44 | 2 | 194 |
| 15 | Thalamus | 23 | -6 | -16 | -6 | 143 |
| 16 | Cuneus/ Precuneus | 7/18/31 | -12 | -73 | 30 | 225 |

Coordinates are in MNI space and correspond to center of the searchlight.

**Table 2.** Reliable clusters in similarity analysis.

| Region | Brodmann<br>areas | x | y | z | Extent<br>(voxels) |
| --- | --- | --- | --- | --- | --- |
| Animals group - domesticity |  |  |  |  |  |
| MFG and dACC | 9/32 | 6.0 | -44.1 | 26.0 | 249 |
| Fruits group - taste |  |  |  |  |  |
| Right IPL and AG | 40 | 57.0 | -57.9 | 34.0 | 85 |

Clusters reliable at $P < 0.05$ corrected (for animals group), $P < 0.02$ corrected (for fruits group). Cluster size thresholded at $P < 0.0001$ . Coordinates are in MNI space and correspond to center of mass of the cluster.

These results suggest that, in a statistically significant number of participants, neural representations of objects (at least when attending to domesticity and taste) reflect behaviorally determined, context-specific similarity structure.

### Connectivity searchlight analysis

We used connectivity analysis to identify areas of activity that were reliably related to the context-specific similarity relationships observed above. To do so, we used areas exhibiting the strongest correlation between neural and behavioral similarity measures as seed regions (Table 1), and computed the correlations between these and the rest of the brain during domesticity trials in the animals experiment, and during taste trials in the fruits experiment (Figure 3a). We used all of the participants’ data to maximize power, and accommodate the possibility that meaningful similarity structure may have been present in some subjects that fell below the stringent level used to determine statistical significance in the foregoing analysis, but may nevertheless be modulated by the activity of areas we sought to identify in the correlational analysis.

For computing the correlations, we used the spatial variance over voxels within the seed region rather than mean activity. Our rationale was that if attention acts to modulate the expression of neural representations by regulating their gain (McAdams and Maunsell, 1999; Treue and Trujillo, 1999; Reynolds et al., 2000; Maunsell and Treue, 2006; Aboitiz and Cosmelli, 2008), then higher gain would be associated with greater spatial variance. This measure also has the virtue of being sensitive to the multivariate nature of the representations (and the similarity structure among them) used to identify the seed regions in the first phase of analysis. Thus, for each seed region and each dimension, we constructed a time series the elements of which were the spatial variance across the voxels in that region at each point in time during which judgments were being made along that dimension (Figure 3b). We then conducted a search for voxels whose activation time series best correlated with the spatial variance time series of the seed region during trials involving each dimension. To make sure that the correlation was dimension-specific, we computed the difference between this correlation and the correlation between the voxel and the seed region during trials of the other dimension (Figure 3a). We then tested the significance of this difference in correlation using a permutation test. Finally, we submitted the number of participants for whom one or more regions exhibited a significant correlation to a binomial test for group level statistics. We found a significant number of participants passed this test for domesticity in the animal group (13 out of 24, binomial *p* (one-tailed) = 10^-6^, average r = 0.31) and taste in the fruit group (9 out of 16, binomial *p* (one-tailed) = 10^-6^, average r = 0.32). In addition, we compared the group average to the null distribution generated by the permutation test, and found the group average to be significant in both domesticity in animals (p = 0.001) and taste in fruits (p = 0.001).

To determine whether using spatial variance provided information that was different from the use of mean activity, we repeated the same analysis using the time series of mean activity instead of spatial variance in the seed regions. We did not find significant results for this analysis in either group (domesticity: 2 out of 24, binomial p (one-tailed) = 0.22, taste: 0 out of 16, binomial p (one-tailed) = 0.44).

To identify the regions that exhibited significant effects in the connectivity analysis at the group level, we used a permutation test to threshold the *p* value and a randomization test to threshold the cluster size. For animals, the largest cluster most significant across participants spanned right inferior frontal gyrus (IFG) and right dlPFC (Brodmann areas 10, 46) for domesticity (p < 0.05 corrected; Figure 3c; Table 3). For fruits, the largest significant cluster spanned left IFG (Brodmann areas 45, 47) and left dlPFC (Brodmann area 10) (p < 0.02 corrected; Figure 3c; Table 3) for taste. Again, repeating the analysis using activation instead of spatial variance did not yield any significant clusters.

**Table 3.**
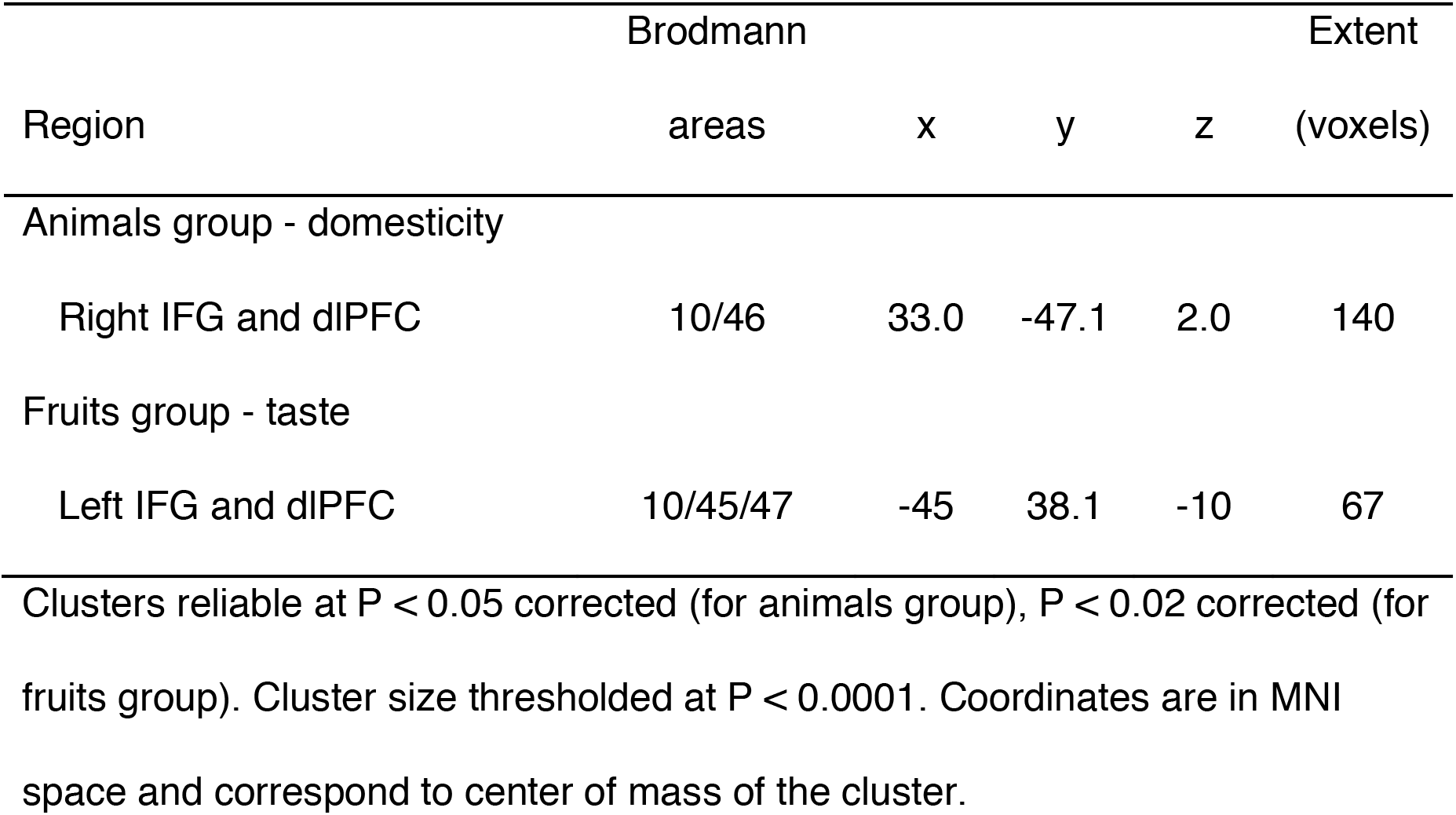
Reliable clusters in connectivity analysis.

## Discussion

We investigated the relationship between perceived similarity relationships among objects and their representations in the brain, and how these are influenced by the direction of attention to different feature dimensions. Specifically, we tested the hypotheses that behavioral judgments of similarity are significantly associated with the similarity of neural representations, and that this relationship is sensitive to the dimension of the objects being attended (e.g., domesticity for animals, or taste for fruits). In support of these hypotheses, we found a significant association between dimension-specific behavioral similarity judgments and neural representations of objects using representational similarity analysis, and found that the expression of these representations was significantly correlated with context-specific activations in the frontal lobe. The results for animals were replicated for fruits in a separate group of participants.

We found clusters in right dlPFC and right IFG that correlated with the expression of similarity along the domesticity dimension in animals, and clusters in left dlPFC and left IFG for taste in fruits. An extensive body of evidence in both human and non-human primates research has associated these regions with the engagement of cognitive control, attention (Duncan, 2001; Miller and Cohen, 2001), and semantic selection (Thompson-Schill et al., 1997). In particular, dlPFC has been shown to be involved in active maintenance of task-specific information or abstract rules versus stimulus identity (Frith and Dolan, 1996; Braver et al., 1997; Cohen et al., 1997; Asaad et al., 2000; MacDonald, 2000; Wallis et al., 2001; Sakai and Passingham, 2003). These findings are broadly consistent with our interpretation that areas in the frontal cortex modulate object representation in a context-specific way to match task demands, and that this in turn influences similarity judgments.

The connectivity analyses yielded significant results using the spatial variance but not the mean activity of the seed regions. Our use of spatial variance as a measure of representational strength was motivated by the idea that attention acts by modulating the expression of representations involved in task performance. This account is consistent with early psychological theories of attention (Treisman, 1969; Treisman and Riley, 1969) and computational modeling of attentional tasks (Cohen et al., 1990). The latter aligns with neurophysiological evidence that attention acts to modulate the sensitivity, or gain of neural processing mechanisms (McAdams and Maunsell, 1999; Treue and Trujillo, 1999; Reynolds et al., 2000; Maunsell and Treue, 2006; Aboitiz and Cosmelli, 2008); that is, excited neurons become more excited and inhibited neurons become less active (Eldar et al., 2013a). We predicted that this effect would be expressed, at the level of patterns of activity in fMRI data, as a higher contrast for attended representations: active voxels within the pattern would be more highly activated, and suppressed voxels would be less active than usual (Eldar et al., 2013b). This, in turn, would produce greater variance in the activity across the voxels within the pattern that we indexed as spatial variance. Therefore, we reasoned that spatial variance should provide a sensitive measure of the expression and attentional modulation of multivariate patterns of activity. We used this to index the expression of dimension-specific neural representations of objects (identified by the correspondence of their similarity structure to behavioral similarity ratings), and then found that this expression was correlated with activity in dlPFC and IFG, which are thought to be involved in task representation, attentional selection and cognitive control (Frith and Dolan, 1996; Braver et al., 1997; Cohen et al., 1997; Asaad et al., 2000; MacDonald, 2000; Duncan, 2001; Miller and Cohen, 2001; Wallis et al., 2001; Sakai and Passingham, 2003).

One important question is why we did not observe similarity effects for size using either animals or fruits. One possibility has to do with the relative nature of size comparisons, and the invariance of object representations to size. For example, the mental representations of the objects may have been “rescaled” based on the particular comparison being made, thus dissociating cardinal representations of size from the individual objects (Frandsen and Holder, 1969; Bundesen and Larsen, 1975; Larsen and Bundesen, 1978; Yamins et al., 2014).

In summary, we showed that the similarity of representations in the brain is modulated by task instructions, such that the way an object is represented reflects its features that are relevant to the task at hand. Furthermore, we showed that the expression of these task-specific representations was correlated with activity in regions of frontal cortex widely thought to be responsible for attentional modulation and cognitive control. We did so using a novel index of representational expression and attention modulation — the spatial variance of patterns of activity identified using MVPA. These findings also provide validation of the usefulness of multivariate pattern and representation similarity analysis for identifying patterns of neural activity associated with mental representations. These methods together with the present findings provide a basis for future work concerning the dynamics of attentional modulation.

## Author Contributions

Formal Analysis, Investigation, Writing – Original Draft, Visualization, W.K.; Supervision, J.D.C and D.O.; Funding Acquisition, J.D.C.; Conceptualization, Methodology, Writing – Review & Editing, all authors

## Acknowledgements

This work is supported by Templeton Foundation. The authors declare no competing financial interests.

